# Genome sequence and annotation of ovine herpesvirus-1

**DOI:** 10.1101/2025.09.04.674317

**Authors:** Paolo Ribeca, Patricia Dewar, Michelle L. McNab, Chris Cousens, George C. Russell, David J. Griffiths

## Abstract

Ovine herpesvirus 1 (OvHV-1) was first identified over 50 years ago in sheep with ovine pulmonary adenocarcinoma (OPA). An aetiological role in OPA was later ruled out and OvHV-1 was found to be a common infection in sheep in several countries. Here, we report the sequence and annotation of the complete OvHV-1 genome. The virus has a similar genomic architecture to members of the *Macavirus* genus of the subfamily *Gammaherpesvirinae* and is most closely related to bovine gammaherpesvirus 6 (BoGHV6). The OvHV-1 genome comprises a 144,637 base pair unique region predicted to encode at least 74 proteins bounded by multiple copies of a 699 base pair GC-rich repetitive terminal repeat. Predicted genes include 61 ORFs conserved among all gammaherpesviruses, and 12 genes present only in macavirus genomes, including a homologue of ovine interleukin-10, previously reported only in ovine gammaherpesvirus-2, and an ornithine decarboxylase, previously described only in BoGHV6. A further gene appears unique to OvHV-1 among macaviruses, encoding a viral-FLIP (FLICE-like inhibitory protein), similar to those found in some other gammaherpesviruses. Notably, several macavirus genes previously predicted in BoGHV6 are defective in OvHV-1. The availability of the genome sequence of OvHV-1 will facilitate studies on its relationship to other macaviruses and its role, if any, in disease.

## INTRODUCTION

Ovine herpesvirus-1 (OvHV-1) was first described over 50 years ago in sheep affected with ovine pulmonary adenocarcinoma (OPA, also known as jaagsiekte), a common neoplastic disease of sheep [1, 2]. Isolates of OvHV-1 were described from the United Kingdom [2], South Africa [3], Kenya [4] and the former Yugoslavia [5], obtained from cultures of macrophages from OPA-affected lung tissue and occasionally from bronchial and mediastinal lymph nodes. Those studies indicated that the virus replicated in cultured ovine alveolar macrophages [1]. Some, but not all, isolates could be transferred to other primary ovine cells [3, 6], although in general the virus was reported to be highly cell-associated and difficult to passage in culture [1].

For several years, OvHV-1 was regarded as a candidate aetiological factor for OPA [7–9], but later studies demonstrated that OPA is instead caused by jaagsiekte sheep retrovirus (JSRV) [10–12], and a direct causative role for OvHV-1 was ruled out. Experimental administration of OvHV-1 to neonatal lambs by intra-tracheal delivery resulted instead in a subclinical interstitial pneumonia [13, 14], but it is unknown whether this pathology is also associated with natural OvHV-1 infection. Although most OvHV-1 isolates were identified in sheep with OPA, serological studies indicated that the virus is a common infection in sheep [15, 16]. This led to the suggestion that OvHV-1 is typically latent in sheep but may be re-activated in some circumstances, such as in the tumour microenvironment of the OPA-affected lung [13]. However, it is unclear whether OvHV-1 can potentiate the pathogenesis of OPA or cause any other disease in sheep.

Following its exclusion as a causative agent for OPA, there has been little new information published on OvHV-1 or its relationship to other ruminant herpesviruses. Two reports have described short genome sequence fragments of a Slovakian isolate of OvHV-1, obtained from choroid plexus cells of a healthy lamb, and comprising partial sequences of capsid proteins ORF25 and ORF26 [6], thymidine kinase (ORF21) and glycoprotein H (ORF22) [17]. PCR with primers specific for these fragments confirmed the similarity of the Slovakian virus with isolates from Scotland and South Africa [6], and established that OvHV-1 is a gammaherpesvirus, most closely related to members of the *Macavirus* genus.

Macaviruses form a genus of gammaherpesviruses that infect various domestic and wild species of ruminants and pigs, and currently nine viral species are formally recognized by the International Committee for Virus Taxonomy (ICTV) [18]. They can be divided into two groups, based on whether they cause malignant catarrhal fever (MCF). MCF-associated viruses include ovine gammaherpesvirus-2 (OvGHV2), alcelaphine gammaherpesviruses-1 and -2 (AlGHV1, AlGHV2), caprine gammaherpesvirus-2 (CpGHV2) and hippotragine gammaherpesvirus, whereas non-MCF-associated macaviruses include bovine gammaherpesvirus-6 (BoGHV6) [19] and suid gammaherpesviruses-5 -6, and -7 (SuGHV5-7) [20, 21]. A pathogenic MCF-virus has also been reported in white-tailed deer [22, 23], and PCR fragments representing a number of related viruses have been described in various farmed ruminants and zoological collections [24–28]. However, these unclassified viruses remain relatively uncharacterised and their association with MCF is unknown.

MCF is a fatal lymphoproliferative disease affecting a variety of artiodactyl species [29]. MCF-associated viruses typically cause a subclinical infection in their adapted host (reservoir) species but cause MCF following cross-species infection into other, non-adapted, hosts. For example, OvGHV2 is enzootic in farmed sheep but causes sporadic cases of MCF if transmitted to cattle. No association of OvHV-1 with MCF has been described but its status within the Macavirus genus and its host range remain to be clarified.

To better establish the relationship of OvHV-1 to other macaviruses, in this study we determined the complete genome sequence of OvHV-1. Phylogenetic analysis found that OvHV-1 clusters with the non-MCF group and is most closely related to BoGHV6 although it also shares some similarities with OvGHV2, an MCF-associated virus. The availability of the complete OvHV-1 genome sequence will permit improved tests for determining the prevalence of infection in sheep and other ruminants and facilitate studies on potential disease associations.

## MATERIALS AND METHODS

### Cells and virus preparation

DNA was extracted from tissue samples and cultured cells with the Qiagen DNeasy blood and tissue kit, as recommended. For preliminary studies, samples of OvHV-1 isolate JSV32 were obtained from a frozen archive held at the Moredun Research Institute. To obtain fresh preparations of OvHV-1 for genome sequencing, cells were isolated from lung lavages from sheep that were naturally clinically affected with OPA. Sheep were euthanised and lungs collected. A sterile stainless-steel funnel was inserted into the trachea and 2 litres of Hank’s balanced salt solution (HBSS) was poured into the lungs. The lungs were massaged gently and the fluid poured out of the lungs into collection vessels through muslin. Cells were collected by centrifugation at 110 × *g* for 20 mins at 4 °C and suspended in HBSS at a concentration of 0.75 ml packed cell volume per 1 ml HBSS, before storing at −80 °C for later use. One of these preparations (case JA548) was subsequently thawed and cultured overnight in four T75 standard tissue-culture flasks at 37 °C, 5% CO_2_ in Dulbecco’s modified Eagle’s medium (Sigma), supplemented with 10% fetal bovine serum and 2 mM glutamine. Adherent cells were then washed once with phosphate buffered saline (PBS) pH 7.4 and cells were lysed by three rounds of freeze-thawing (−80 °C and 20 °C; 25 ml culture medium per flask). The supernatant from the lysed cells was then harvested and clarified by centrifugation (870 × *g*, 4 °C, 20 mins), before pelleting virus particles by ultracentrifugation (100,000 × *g*, 4 °C, 90 mins). The virus pellet was suspended in 100 µl TNE (10 mM Tris-HCl, 100 mM sodium chloride, 1 mM EDTA; pH 7.8) and DNA extracted and purified (Qiagen DNeasy). From this, we obtained approximately 30 µg of DNA, which was sequenced using Illumina short-read sequencing. Additional DNA samples were prepared from lung tissue (case JA542) and lung lavage cells (cases JA502, JA555 and JA564) obtained from OPA-affected sheep from farms located in Scotland or Northern England.

### Ethics statement

All studies involving sheep were performed with local and national ethical approval in accordance with the Animal (Scientific Procedures) Act 1986, under Home Office licence PPL 60/3870.

### Polymerase chain reaction

Degenerate primer PCR using ‘pan-herpesvirus’ primers [30] was performed with a mixture of degenerate oligonucleotide primers using Taq polymerase (Promega). Additional reactions with primers specific for OvHV-1 were performed with KOD Hot Start DNA polymerase (Merck) as recommended. All primer sequences are shown in Table 1.

**Table 1.**
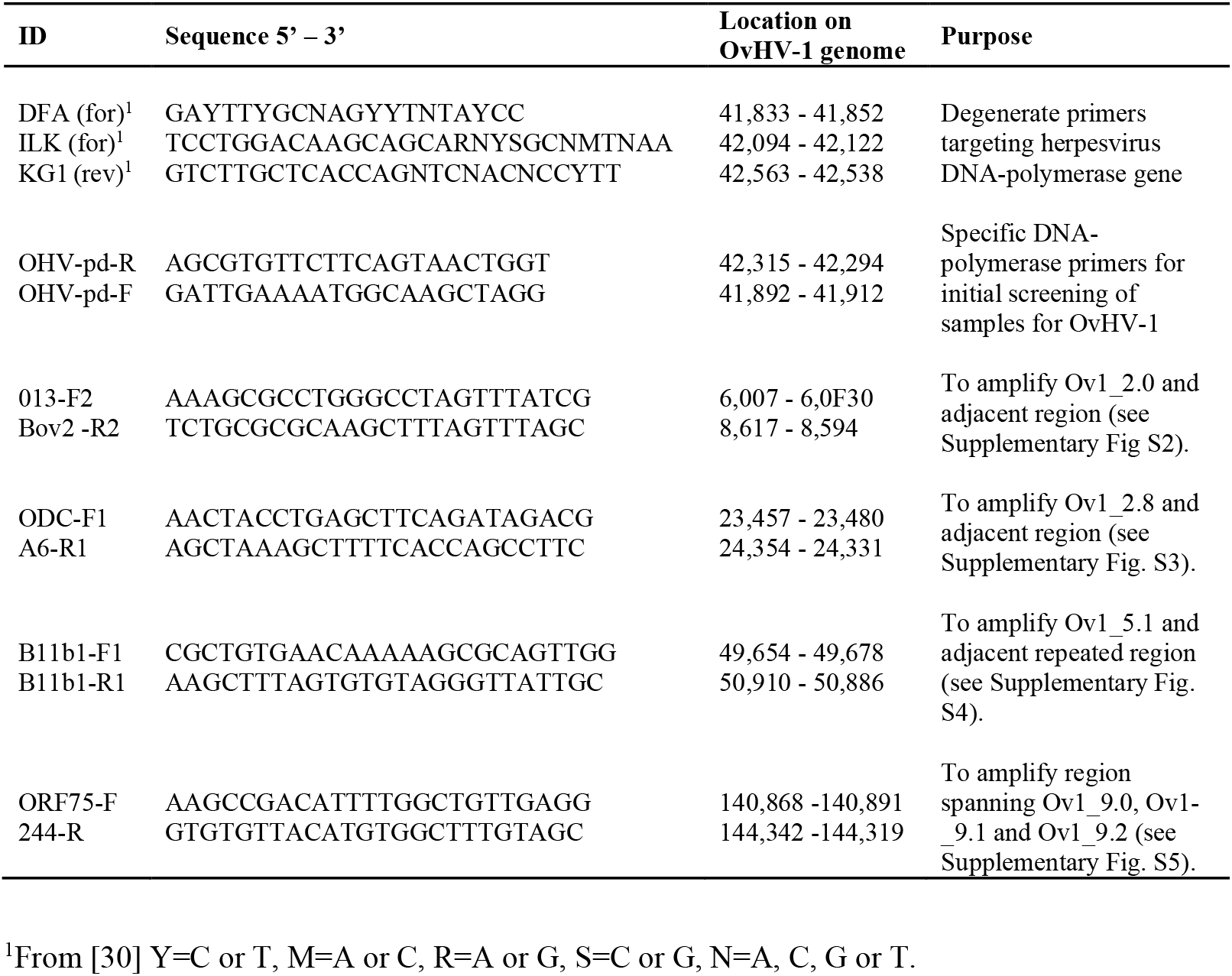
Oligonucleotides used in this study.

### DNA Sequencing

DNA sequencing libraries were prepared using an Illumina TruSeq DNA library kit and sequencing was performed on an Illumina HiSeq2000 sequencing instrument yielding 101 base-pair paired end reads. Note that the sequencing was performed in 2010, using an early generation Illumina base-calling pipeline in which each read retained the adapter sequence. Here, the adapters were removed by manual deletion of the first 6 bases of each read. This is essential to obtain a high-quality assembly. Sanger sequencing of PCR products was performed directly on purified amplicons using internal primers (Eurofins Genomics) and analysed using DNASTAR Lasergene software. Clustal Omega alignments were performed with the EMBL-EBI Job Dispatcher sequence analysis tools framework [31]. Protein motif searches were performed using the Prosite database (https://prosite.expasy.org/) [32].

### Genome assembly and annotation

Following adapter removal, genome assembly was performed with NINJA, a bioinformatics pipeline based on a novel overlap-layout-consensus method that will be described in a separate manuscript. Briefly, NINJA provides specialised facilities to reconstruct microbial genomes, such as optimised linearization of the assembly graph, realignment-based polishing of short genomic features, determination of repeat copy number, and variant calling based on a zero-ploidy low-frequency model [33]. NINJA was run with standard parameters, except that a minimum coverage of 3 for contig extension was required.

The quality of the final draft genome was checked by realigning reads to the draft assembly to check that no misassembly was evident. Reads were mapped to the draft OvHV-1 genome sequence and to the sheep genome (ARS-UI_Ramb_v2.0, INSDC Assembly GCA_016772045.1) [34] with the GEM mapper [35] by keeping the best stratum of alignments (i.e., the set of best alignments with minimum edit distance) and, wherever multiple alignments were present, by randomly selecting one of them to keep the number of reads constant. Assuming uniform sequencing depth across the genome, this procedure also allowed estimation of the number of copies in the terminal repeats as the ratio between the coverage of repeats and the average coverage of the genome.

To directly compare OvHV-1 and BoGHV6 genomes by dot plot analysis, the two genome sequences were split into consecutive sequence fragments of 80 nucleotides, with each fragment overlapping the previous one by 40 nucleotides. Each fragment from one genome was then aligned to the other genome with BLASTN [36]. All the alignments of sufficient quality (sequence similarity ≥ 90%) were plotted as dots at their respective positions within each sequence.

The draft OvHV-1 genome sequence was then annotated with The Transporter [37], a command-line reimplementation of GATU [38] that has been used previously for other large DNA viruses [39]. Briefly, the method extracts features from a specified annotation (or set of annotations) and builds a database. Here, the input database of features was built from the annotations of an extensive collection of gammaherpesvirus reference genomes downloaded from NCBI. Features were then searched for in the OvHV-1 genome using several methods including splign [40] to identify spliced genes, BLASTP, TBLASTN, and transeq [31] to identify additional ORFs.

Where several overlapping transported features are found at the same location on the target genome, they were manually reviewed to select the most biologically relevant, based on Transporter ranking.

### Phylogenetic analysis and synteny

Phylogenetic analysis of gammaherpesvirus proteins was performed by extracting all annotated proteins from the GenBank file of each virus and indexing in a BLAST database. The OvHV-1 proteins in the region from ORF19 to ORF45, which are highly conserved across all the viruses considered, were then used to query the database. The orthologues thus identified for each genome were then concatenated in order and the resulting sequences were aligned with MAFFT version 7.526 [41, 42]. The resulting multiple alignment was provided to IQtree [43] to perform model optimisation at standard parameters and bootstrapping with 1000 replicates. The resulting consensus tree was essentially identical to a tree generated with FastTree [44] for comparison purposes. Homologous gene clusters in gammaherpesvirus genomes were visualised using clinker [45].

### Accession numbers

The Illumina sequencing data have been deposited in the GenBank Short Read Archive under accession number PRJNA1268940. The assembled OvHV-1 genome sequence has GenBank accession number PV694339.

## RESULTS AND DISCUSSION

### Identification of a fragment of OvHV-1 DNA polymerase sequence by degenerate-primer PCR

In initial experiments, a 685 bp fragment of the OvHV-1 genome sequence was obtained from a virus stock archived at the Moredun Research Institute in the 1980s (isolate JSV32) using degenerate-primer PCR targeting a conserved region of the DNA polymerase gene [30].

Sequencing of the PCR product identified a sequence with up to 80% nucleotide similarity with macaviruses. Specific PCR primers were then designed from this sequence for detection of OvHV-1 in other samples (Table 1).

### Genome sequencing of OvHV-1

To acquire new material for virus isolation and genome sequencing, cells were obtained from lung lavages of sheep that were clinically affected with OPA. The cells were allowed to adhere to plastic in tissue culture flasks overnight before harvesting, isolating DNA and testing for OvHV-1 DNA by specific PCR for the DNA-polymerase region (data not shown). One positive sample (sheep JA548) was further processed to generate DNA for genome sequencing. Consistent with earlier reports [1], OvHV-1 was found to be cell-associated, therefore a freeze-thaw procedure was used to generate soluble cell extracts, from which virus particles were concentrated by ultracentrifugation. DNA extracted from this material was sequenced using Illumina short read sequencing.

Sequencing yielded 6,425,055 read-pairs of length 2 × 101 bases (2 × 95 bases after adaptor removal). Of these, 5,605,679 reads (87.2%) mapped to the sheep reference genome. The remaining reads were assembled *de novo* using a custom pipeline (see Materials and Methods for a detailed description), which produced a complete OvHV-1 draft genome sequence of 146,035 bp that includes terminal repeats. Following assembly, alignment of total sequencing data resulted in 154,315 paired end reads (2.4%) mapping to the final genome assembly, with an average coverage of approximately 200× (Fig. 1). There were no coverage gaps and the only dips observed corresponded to repetitive regions which are challenging to sequence, as commonly observed in comparable viruses.

**Figure 1.**
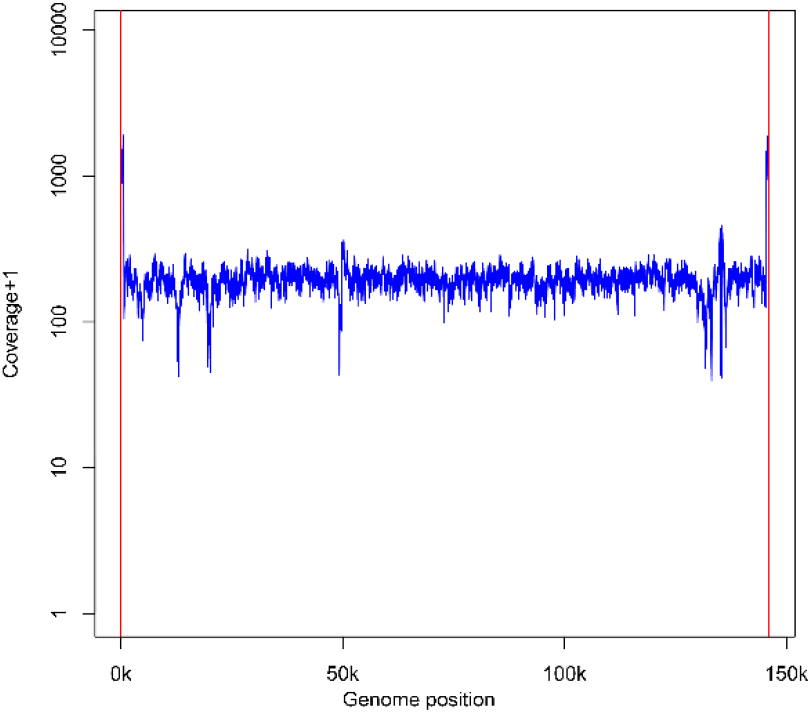
–Coverage plot of sequencing reads against the OvHV-1 reference genome. Following *de novo* genome assembly, all reads were mapped back to the reference genome. Note the vertical axis is in logarithmic scale.

Consistent with other gammaherpesviruses, the OvHV-1 genome comprises a GC-poor L-DNA unique region (38.8% GC) that encodes the structural and non-structural viral proteins, bounded at both ends by GC-rich repetitive terminal H-DNA regions (65% GC). Notably, the sequence of this Scottish isolate has 99% nucleotide identity with regions of the OvHV-1 genome described previously from a Slovakian isolate [6, 17], confirming this as OvHV-1. Fig. 2 is a representation of the assembled genome with key annotated features marked.

**Figure 2.**
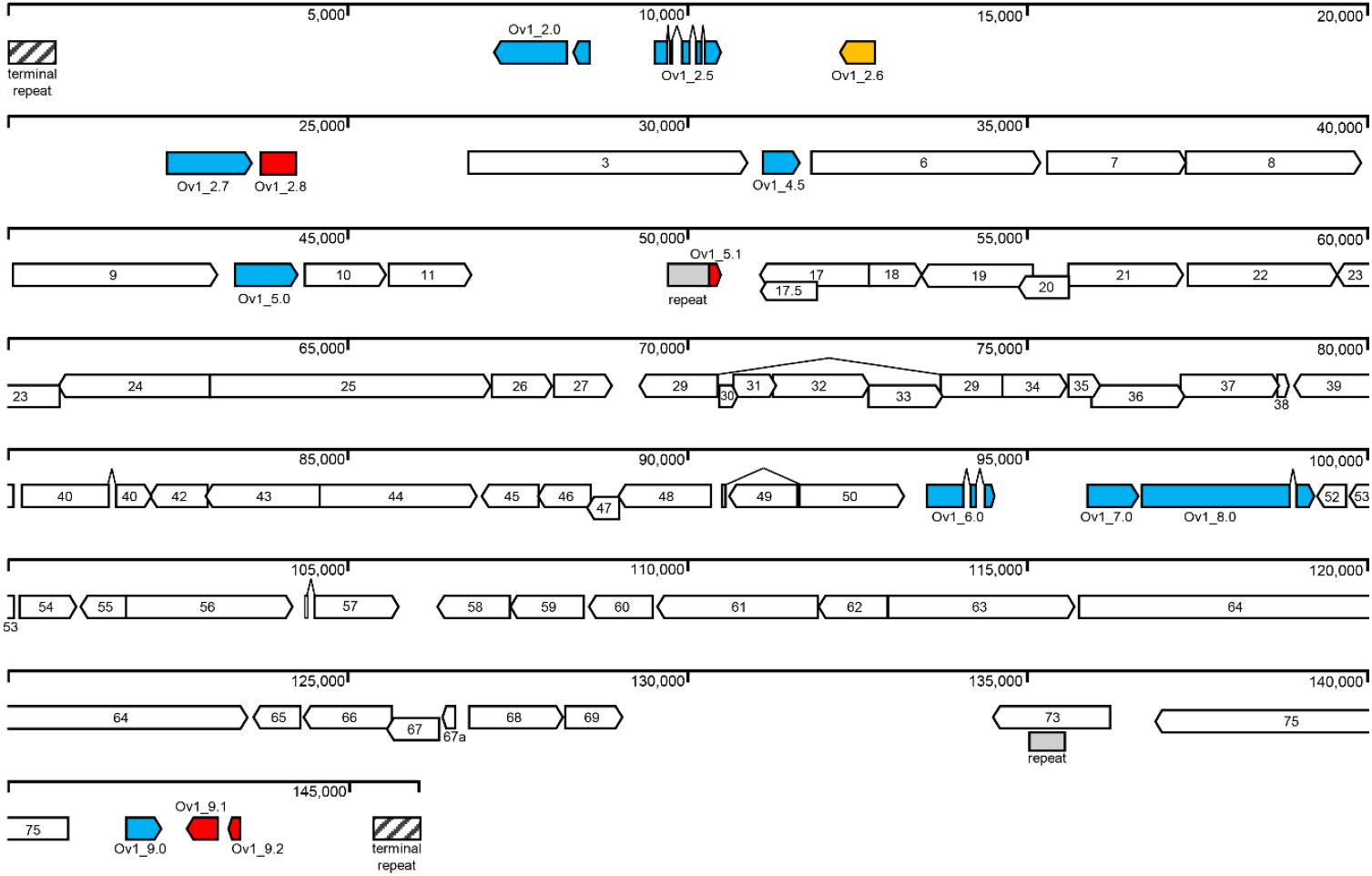
Organisation of the OvHV-1 genome. The figure shows the organisation of predicted protein-coding regions of the OvHV-1 genome (see also Table 2). White arrows indicate conserved gammaherpesvirus ORFs with numbers based on the annotation of SaGHV2 [46]. Blue arrows indicate intact Macavirus-specific genes. The yellow arrow indicates an ORF predicted to encode a macavirus vFLIP-like protein. Red arrows indicate truncated or defective homologues of BoGHV6 annotated proteins.

### Repeat regions

The terminal direct repeats have a unit length of 699 nucleotides and one copy of the terminal repeat was included at each end of the reference genome sequence in the GenBank entry. Based on the assumption of uniform sequencing depth, normalised alignment of sequencing reads to the genome resulted in higher coverage observed in the repeats (Fig. 1). Estimation of coverage ratios indicated approximately 12 to 16 repeat copies per genome (i.e., 6 to 8 copies in each of the 5′ and 3′ terminal repeats) in the purified cell-associated virus. In addition, and consistent with the structure of viruses belonging to the same genus, several internal repeats were identified, which are summarised in Supplementary Table S1. The two largest regions are at nucleotide positions 49,764 (close to Ov1_5.1) and 135,145 (within ORF73). The copy number of these longer repeats was estimated from short-read coverage and complemented with Sanger sequencing because the total length of each tandem repeat region was greater than the sequencing read length. The region within ORF73 is G-poor and hypervariable, with a complex sequence of pseudo-repeats that are likely to differ from isolate to isolate. This region is predicted to encode an extended acidic repeat in the ORF73 protein, in common with other gammaherpesviruses which also encode long charged regions in this protein [19, 46–48].

**Table 2.** Predicted proteins encoded by OvHV-1.

| Annotation (exon) | Protein Product Name | CDS Start | CDS end | Length (nt) | Strand orientation | OvHV-1 vs. closest BoGHV-6 homologue |  |  |
| --- | --- | --- | --- | --- | --- | --- | --- | --- |
|  |  |  |  |  |  | Identities | positives | gaps |
| Ov1_2.0 (1) | Bov1.b2, basic leucine zipper motif | 8575 | 8357 | 219 | R | 318/440(72%) | 355/440(80%) | 21/440(4%) |
| Ov1_2.0 (2) |  | 8258 | 7182 | 1077 |  |  |  |  |
| Ov1_2.5 (1) | Interleukin-10 | 9535 | 9687 | 153 | F | 12/20(60%) | 17/20(85%) | 0/20(0%) |
| Ov1_2.5 (2) |  | 9764 | 9823 | 60 |  |  |  |  |
| Ov1_2.5 (3) |  | 9908 | 10060 | 153 |  |  |  |  |
| Ov1_2.5 (4) |  | 10141 | 10206 | 66 |  |  |  |  |
| Ov1_2.5 (5) |  | 10433 | 10522 | 90 |  |  |  |  |
| Ov1_2.6 | CFLAR/vFLIP | 12741 | 12256 | 486 | R | not present |  |  |
| Ov1_2.7 | ornithine decarboxylase | 22373 | 23662 | 1290 | F | 338/431(78%) | 390/431(90%) | 2/431(0%) |
| Ov1_2.8 | Bov2.b3 (truncated) | 24291 | 23226 | 66 | R | 9/21 (43%) | 12/21(57%) | 0/20(0%) |
| ORF3 | tegument protein/v-FGAM-synthetase | 26799 | 30941 | 4143 | F | 1132/1378(82%) | 1262/1378(91%) | 0/1378(0%) |
| Ov1_4.5 | vBcl-2 | 31124 | 31666 | 543 | F | 133/182(73%) | 159/182(87%) | 2/182(1%) |
| ORF6 | single-stranded DNA binding protein | 31824 | 35222 | 3399 | F | 1027/1133(91%) | 1091/1133(96%) | 1/1133(0%) |
| ORF7 | terminase subunit | 35275 | 37323 | 2049 | F | 549/683(80%) | 601/683(87%) | 1/683(0%) |
| ORF8 | glycoprotein B | 37316 | 39880 | 2565 | F | 741/861(86%) | 797/861(92%) | 14/861(1%) |
| ORF9 | DNA polymerase | 40111 | 43104 | 2994 | F | 890/999(89%) | 938/999(93%) | 4/999(0%) |
| Ov1_5.0 | G protein-coupled receptor | 43384 | 44331 | 948 | F | 236/296(80%) | 263/296(88%) | 0/296(0%) |
| ORF10 | ORF10 | 44388 | 45596 | 1209 | F | 298/403(74%) | 340/403(84%) | 1/403(0%) |
| ORF11 | ORF11 | 45618 | 46862 | 1245 | F | 339/414(82%) | 371/414(89%) | 0/414(0%) |
| <sup>1</sup> Ov1_5.1 | Bov11.b1 (truncated) | 50354 | 50545 | 192 | F | 37/63(59%) | 43/63(68%) | 3/63(4%) |
| ORF17.5 | capsid scaffold protein | 51915 | 51079 | 837 | R | 194/293(66%) | 222/293(75%) | 24/293(8%) |
| ORF17 | protease; capsid protein | 52710 | 51079 | 1632 | R | 395/551(72%) | 444/551(80%) | 28/551(5%) |
| ORF18 | ORF18 | 52709 | 53497 | 789 | F | 220/259(85%) | 237/259(91%) | 0/259(0%) |
| ORF19 | tegument protein | 55114 | 53471 | 1644 | R | 479/547(88%) | 521/547(95%) | 0/547(0%) |
| ORF20 | nuclear protein | 55655 | 54945 | 711 | R | 205/236(87%) | 221/236(93%) | 0/236(0%) |
| ORF21 | thymidine kinase | 55657 | 57348 | 1692 | F | 443/564(79%) | 495/564(87%) | 3/564(0%) |
| ORF22 | glycoprotein H | 57386 | 59596 | 2211 | F | 636/736(86%) | 695/736(94%) | 0/736(0%) |
| ORF23 | ORF23 | 60798 | 59593 | 1206 | R | 341/401(85%) | 377/401(94%) | 0/401(0%) |
| ORF24 | ORF24 | 63010 | 60776 | 2235 | R | 637/749(85%) | 702/749(93%) | 5/749(0%) |
| ORF25 | major capsid protein | 63012 | 67103 | 4092 | F | 1236/1363(91%) | 1315/1363(96%) | 0/1363(0%) |
| ORF26 | capsid triplex subunit 2 | 67140 | 68054 | 915 | F | 273/304(90%) | 294/304(96%) | 0/304(0%) |
| ORF27 | glycoprotein | 68059 | 68943 | 885 | F | 237/293(81%) | 265/293(90%) | 4/293(1%) |
| ORF29 (1) | DNA packaging terminase subunit | 74645 | 73731 | 915 | R | 612/687(89%) | 650/687(94%) | 0/687(0%) |
| ORF29 (2) |  | 70497 | 69349 | 1149 |  |  |  |  |
| ORF30 | ORF30 | 70512 | 70760 | 249 | F | 61/81(75%) | 73/81(90%) | 0/81(0%) |
| ORF31 | ORF31 | 70685 | 71356 | 672 | F | 179/223(80%) | 202/223(90%) | 0/223(0%) |
| ORF32 | ORF32 | 71305 | 72732 | 1428 | F | 333/481(69%) | 387/481(80%) | 11/481(2%) |
| ORF33 | tegument protein | 72722 | 73741 | 1020 | F | 318/339(94%) | 330/339(97%) | 0/339(0%) |
| ORF34 | ORF34 | 74644 | 75624 | 981 | F | 275/326(84%) | 304/326(93%) | 0/326(0%) |
| ORF35 | ORF35 | 75602 | 76054 | 453 | F | 126/153(82%) | 138/153(90%) | 3/153(1%) |
| ORF36 | putative tyrosine kinase | 75948 | 77333 | 1386 | F | 410/463(89%) | 434/463(93%) | 2/463(0%) |
| ORF37 | alkaline exonuclease | 77266 | 78723 | 1458 | F | 441/485(91%) | 465/485(95%) | 0/485(0%) |
| ORF38 | ORF38 | 78678 | 78866 | 189 | F | 51/63(81%) | 55/63(87%) | 1/63(1%) |
| ORF39 | glycoprotein M | 80059 | 78926 | 1134 | R | 342/377(91%) | 358/377(94%) | 0/377(0%) |
| ORF40 (1) | helicase-primase complex | 80198 | 81486 | 1289 | F | 458/644(71%) | 523/644(81%) | 36/644(5%) |
| ORF40 (2) |  | 81565 | 82108 | 544 |  |  |  |  |
| ORF42 | ORF42 | 82924 | 82091 | 834 | R | 244/277(88%) | 262/277(94%) | 0/277(0%) |
| ORF43 | minor capsid protein | 84569 | 82890 | 1680 | R | 504/557(90%) | 536/557(96%) | 0/557(0%) |
| ORF44 | helicase-primase subunit | 84559 | 86901 | 2343 | F | 711/787(90%) | 750/787(95%) | 7/787(0%) |
| ORF45 | ORF45 | 87785 | 86967 | 819 | R | 200/275(73%) | 227/275(82%) | 5/275(1%) |
| ORF46 | uracil DNA glycosylase | 88566 | 87808 | 759 | R | 229/252(91%) | 239/252(94%) | 0/252(0%) |
| ORF47 | ORF47 | 88978 | 88547 | 432 | R | 103/127(81%) | 112/127(88%) | 0/127(0%) |
| ORF48 | ORF48 | 90351 | 88984 | 1368 | R | 329/464(71%) | 379/464(81%) | 11/464(2%) |
| ORF49 | transcriptional control protein | 91605 | 90616 | 990 | R | 274/329(83%) | 304/329(92%) | 0/329(0%) |
| ORF50 (1) | Rta-transactivator | 90514 | 90567 | 54 | F | 408/504(81%) | 436/504(86%) | 10/504(1%) |
| ORF50 (2) |  | 91639 | 93204 | 1566 |  |  |  |  |
| Ov1_6.0 (1) | Bov6 | 93523 | 94083 | 561 | F | 224/285(79%) | 237/285(83%) | 18/285(6%) |
| Ov1_6.0 (2) |  | 94160 | 94276 | 117 |  |  |  |  |
| Ov1_6.0 (3) |  | 94354 | 94518 | 165 |  |  |  |  |
| Ov1_7.0 | putative glycoprotein | 95901 | 96683 | 783 | F | 208/254(82%) | 228/254(89%) | 0/254(0%) |
| <sup>1</sup> Ov1_8.0 (1) | putative major envelope glycoprotein | 96689 | 98645 | 1957 | F | 397/775(51%) | 502/775(64%) | 52/775(6%) |
| <sup>1</sup> Ov1_8.0 (2) |  | 98968 | 99257 | 290 |  |  |  |  |
| ORF52 | ORF52 | 99701 | 99282 | 420 | R | 126/139(91%) | 135/139(97%) | 1/139(0%) |
| ORF53 | ORF53 | 100088 | 99771 | 318 | R | 81/104(78%) | 87/104(83%) | 0/104(0%) |
| ORF54 | dUTPase | 100176 | 101048 | 873 | F | 255/289(88%) | 271/289(93%) | 0/289(0%) |
| ORF55 | ORF55 | 101751 | 101083 | 669 | R | 198/213(93%) | 207/213(97%) | 0/213(0%) |
| ORF56 | helicase-primase primase subunit | 101750 | 104251 | 2502 | F | 708/835(85%) | 774/835(92%) | 6/835(0%) |
| ORF57 (1) | transcriptional control protein | 104390 | 104441 | 52 | F | 358/459(78%) | 390/459(84%) | 10/459(2%) |
| ORF57 (2) |  | 104519 | 105816 | 1298 |  |  |  |  |
| ORF58 | ORF58 | 107403 | 106348 | 1056 | R | 316/351(90%) | 336/351(95%) | 0/351(0%) |
| ORF59 | processivity factor | 108501 | 107407 | 1095 | R | 325/375(87%) | 340/375(90%) | 11/375(2%) |
| ORF60 | ribonucleotide-reductase, small subunit | 109519 | 108602 | 918 | R | 290/305(95%) | 297/305(97%) | 0/305(0%) |
| ORF61 | ribonucleotide-reductase, large subunit | 111948 | 109588 | 2361 | R | 707/786(90%) | 748/786(95%) | 2/786(0%) |
| ORF62 | capsid triplex subunit 1 | 112991 | 111978 | 1014 | R | 275/339(81%) | 308/339(90%) | 2/339(0%) |
| ORF63 | tegument protein | 112993 | 115776 | 2784 | F | 737/928(79%) | 822/928(88%) | 2/928(0%) |
| ORF64 | large tegument protein | 115819 | 123579 | 7761 | F | 1877/2620(72%) | 2191/2620(83%) | 40/2620(1%) |
| ORF65 | capsid protein | 124286 | 123615 | 672 | R | 151/227(67%) | 168/227(74%) | 28/227(12%) |
| ORF66 | ORF66 | 125643 | 124330 | 1314 | R | 345/437(79%) | 384/437(87%) | 2/437(0%) |
| ORF67 | tegument protein | 126344 | 125553 | 792 | R | 224/267(84%) | 235/267(88%) | 6/267(2%) |
| ORF67a | ORF67a | 126608 | 126354 | 255 | R | 71/84(85%) | 81/84(96%) | 0/84(0%) |
| ORF68 | putative major envelope glycoprotein | 126801 | 128198 | 1398 | F | 378/460(82%) | 423/460(91%) | 0/460(0%) |
| ORF69 | ORF69 | 128200 | 129063 | 864 | F | 257/286(90%) | 273/286(95%) | 0/286(0%) |
| <sup>2</sup> ORF73 | putative immediate early protein | 136223 | 134499 | 2725 | R | 99/159(62%) | 115/159(72%) | 8/159(5%) |
| ORF75 | FGAM-synthase | 140877 | 136927 | 3951 | R | 1066/1317(81%) | 1198/1317(90%) | 3/1317(0%) |
| Ov1_9.0 | vBcl-2 (Bov9) | 141697 | 142224 | 528 | F | 150/174(86%) | 160/174(91%) | 2/174(1%) |
| Ov1_9.1 | Bov9.b1 (truncated) | 143053 | 142613 | 441 | R | 50/143(35%) | 72/143(50%) | 18/143(12%) |
| Ov1_9.2 | Bov9.b2 (truncated) | 143365 | 143207 | 159 | R | 33/48(69%) | 42/48(87%) | 0/48(0%) |

<sup>1</sup> Ov1\_8.0 is annotated as spliced according to its orthologue in BoGHV6 [19]. While the N and C-terminal parts of the proteins show high similarity between the two viruses, the intermediate region is quite variable. Although the length of the second exon appears to be conserved, it is unclear whether that is the case for the first exon and several positions for the splice site would be possible, for instance 96689-98645 or 96689-98882. Here, the first one has been chosen, which gives the first exon a length similar to that in BoGHV6.
<sup>2</sup> For ORF73, the BLASTP analysis is for 162 amino acids at the C-terminus, after the acidic repeat region.

### Protein coding potential of OvHV-1

The OvHV-1 genome sequence contains 74 predicted open reading frames that were annotated with functions based on similarity to previously identified herpesvirus proteins. This includes 61 core proteins that are conserved among gammaherpesviruses. By convention, these have been annotated by ORF number according to their orthologue in *Saimiriine gammaherpesvirus 2* (SaGHV-2; also known as Herpesvirus saimiri [46]; Fig. 2, Table 2). There are also 12 ‘non-core’ predicted proteins shared with other macaviruses and a further protein that is apparently novel to OvHV-1. These have been prefixed “Ov1_” in the genome annotation and suffixed with numbering consistent with the designations used in the genomes of AlGHV1 [49], OvGHV2 [47] and BoGHV6 [19] (Table 3). Note that, for BoGHV6, the annotations in the publication [19] and the GenBank entry (KJ705001) differ, particularly in relation to spliced genes (Bov2, ORF50 and Bov8). For the present report the annotation from the Jia et al (2014) publication [19] was used because that is more consistent with those of AlGHV1 and OvGHV2. Four of the predicted ORFs with similarity to proteins annotated in BoGHV6 are truncated by in-frame stop codons (see below).

**Table 3.**
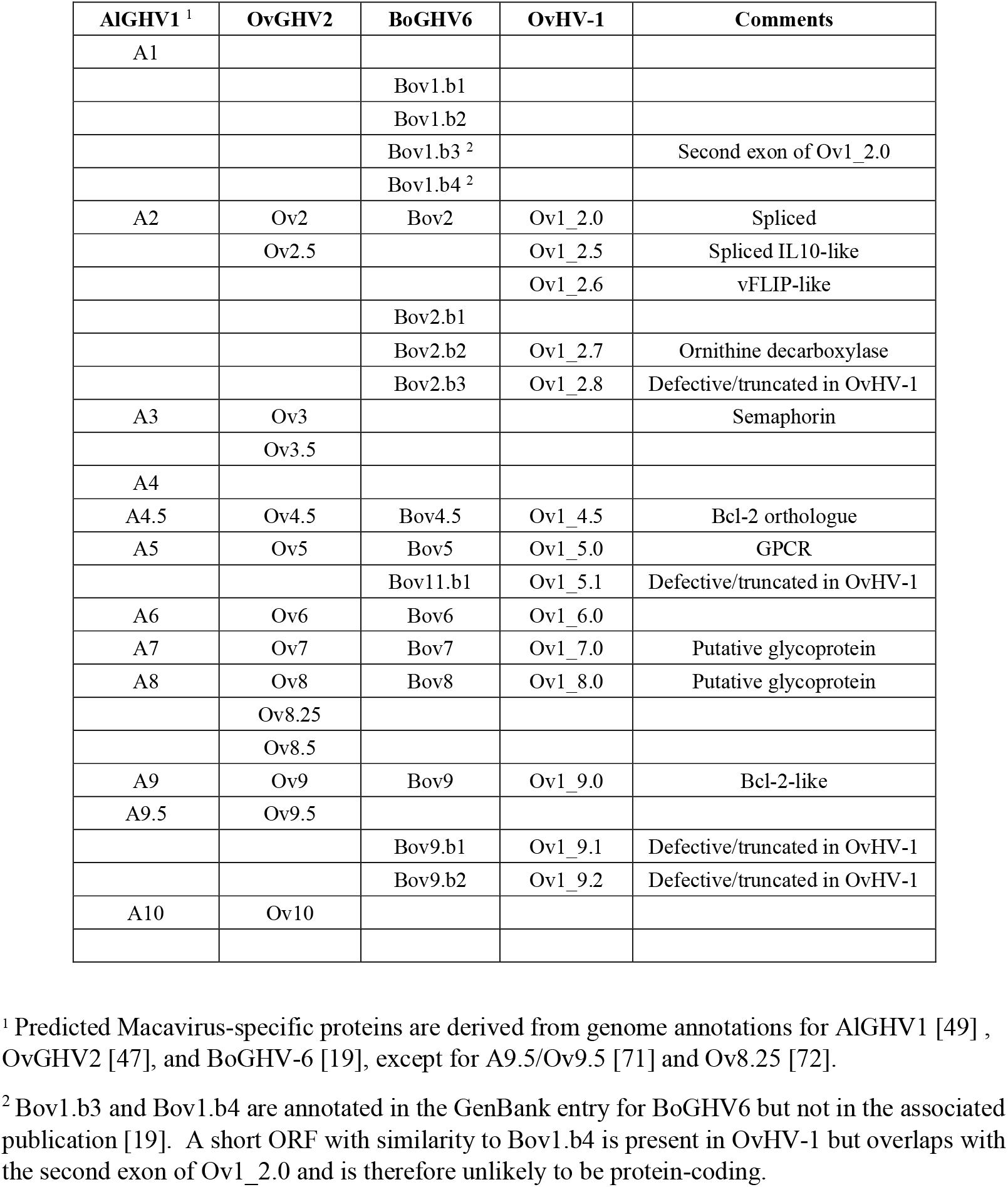
Macavirus-specific proteins.

Comparison of OvHV-1 genome structure with other ruminant gammaherpesviruses indicates a high level of synteny with BoGHV6, as illustrated by dot plot analysis (Fig. 3a) and genome comparison (Supplementary Fig. S1). Among the core genes, OvHV-1 and BoGHV6 share amino acid identity in the range 75%-93%. Other features in common with BoGHV6 are the presence of an extended region at the left-hand end of the genome, which is relatively gene-poor, and the presence of a gene encoding a potential ornithine decarboxylase (Ov1_2.7; see below), a function previously only identified in BoGHV-6 among herpesviruses [19]. Phylogenetic analysis supports clustering of OvHV-1 with BoGHV6 within the Macaviruses (Fig. 3b). Interestingly, these two ruminant non-MCF viruses appear more closely related to the MCF-associated viruses than to the suid macaviruses.

**Figure 3.**
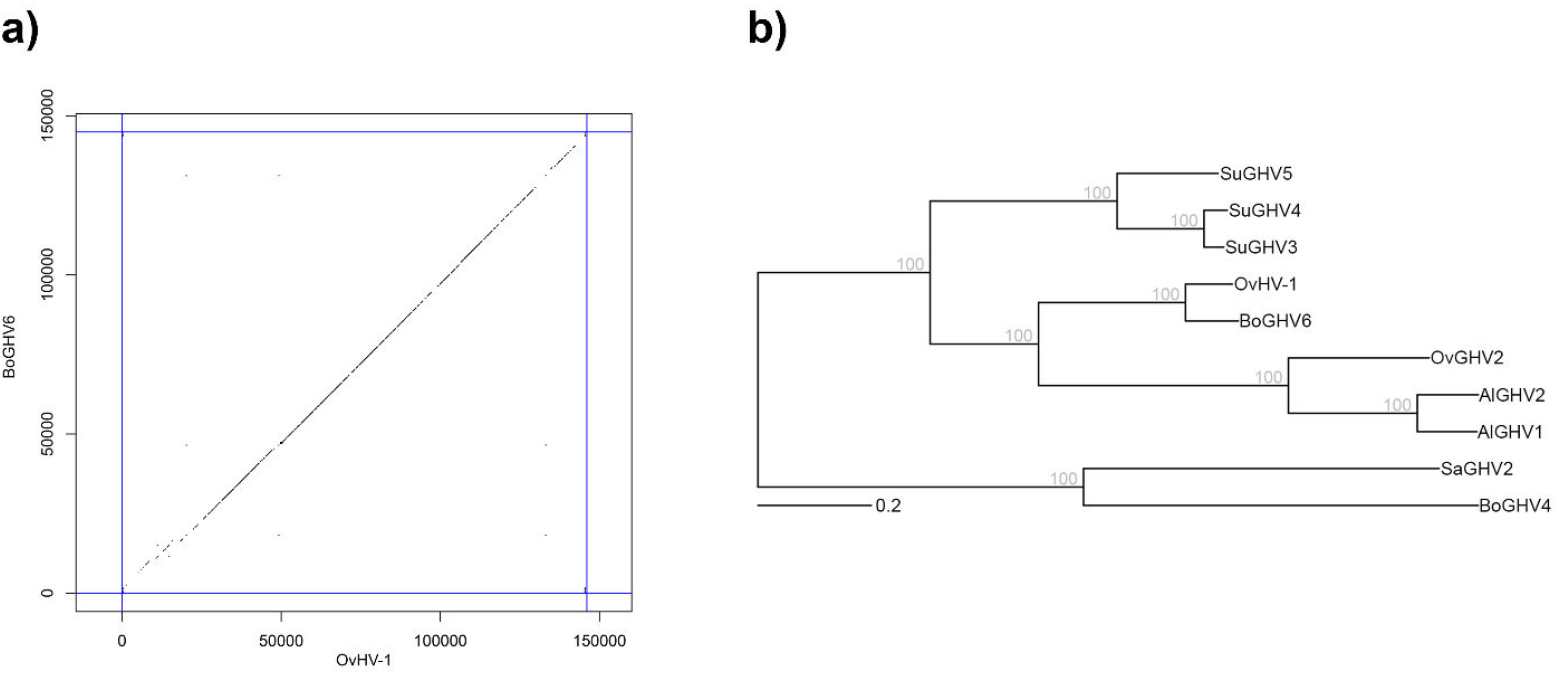
– Phylogenetic analysis of OvHV-1 and Macaviruses. **a)**. Sequence synteny of OvHV-1 with BoGHV6 visualised as a dot plot. Each dot represents a short subsequence of the same length that is present in both sequences. **b)**. Maximum-likelihood tree inferred from concatenated protein sequences (ORF 19 to ORF 45). Bootstrap analysis (1000) indicated 100 % support at all nodes. Bar = estimated changes per amino acid position. GenBank accession codes for each virus sequence are: suid gammaherpesvirus 3 (SuGHV3), AF478169; SuGHV4, AY170317; SuGHV5, AY170316; OvHV-1, PV694339; AlGHV1, AF005370; AlGHV2, KF274499; OvGHV2, AY839756; BoGHV6, KJ705001. Saimiriine gammaherpesvirus 2 (SaGHV2; X64346) and bovine gammaherpesvirus 4 (BoGHV4; AF318573) were included as more distant relatives.

### Macavirus genes in the OvHV-1 genome

Ov1_2.0 is a positional orthologue of AlGHV1 A2 and OvGHV2 Ov2, which are basic leucine zipper proteins related to bZIP proteins found in other gammaherpesviruses and thought to have roles in regulating various processes including host and viral gene transcription and apoptosis [50, 51]. Ov1_2.0 contains an imperfect leucine heptad repeat suggesting it may have a similar function to A2 or Ov2. However, similar to Bov2 of BoGHV6, Ov1_2.0 encodes an extended C-terminal region (Supplementary Fig. S2), which could confer functional differences compared to A2/Ov2. Interestingly, homologues in each of these macaviruses are encoded by two exons, with a splice site as the same position (Supplementary Fig. S2 and Table S2).

OvHV-1 encodes a homologue of ovine interleukin-10 (IL-10), designated Ov1_2.5. Interestingly, an IL-10 gene is also present in OvGHV2, an MCF-associated macavirus, but is absent from BoGHV6 except for a short 20 amino acid fragment that appears unlikely to encode a functional protein (Bov2.5, [19]). Both the OvHV-1 and OvGHV2 homologues are encoded by five exons with a splicing pattern identical to that of the ovine IL-10 gene indicating capture from the host genome, although intronic sequences are not conserved. In addition, OvHV-1 Ov1_2.5 shares greater amino acid identity with ovine IL-10 (84% in the mature protein) than does the OvGHV2 IL-10 (50%; Fig, 4a). There are also IL-10-like proteins encoded by other gammaherpesviruses, including Epstein Barr virus (EBV) and equine herpesvirus-2 (EHV-2) and by parapox viruses, including the ovine parapoxvirus orf [52], although none of those homologues has the splicing pattern of their host. Orf virus IL-10 is also highly similar (81%) to the ovine protein and was reported to be fully functional [53]. Preservation of a functional IL-10 coding sequence indicates an important role for this protein in modulating the immune response in OvHV-1 infection and promoting viral persistence. However, this requirement appears to have been lost for BoGHV6.

**Figure 4.**
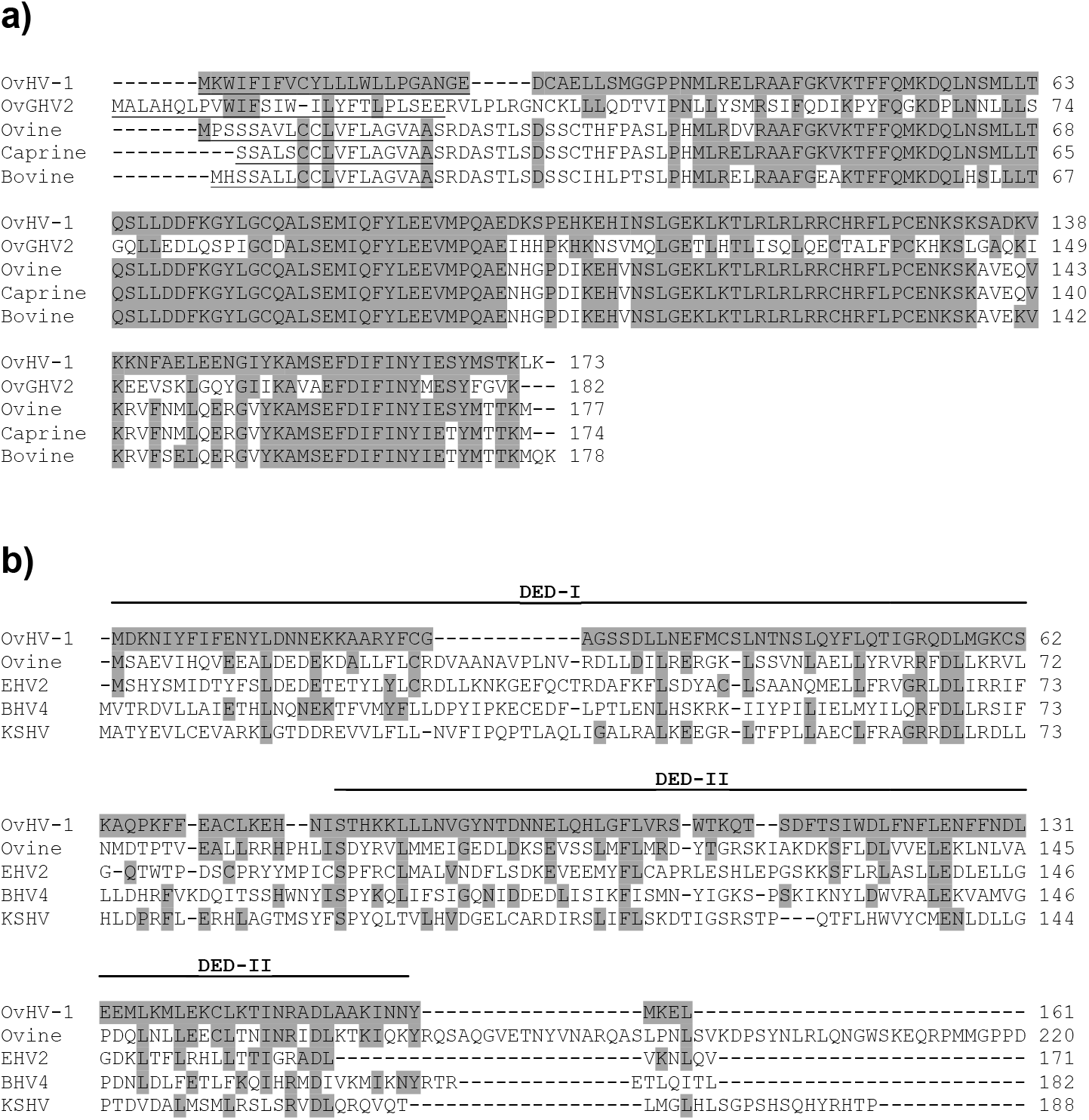
OvHV-1 IL-10 and vFLIP-like protein sequences. The figure shows the predicted protein sequences of (a) IL-10-like (Ov1_2.5) and (b) putative vFLIP (Ov1_2.6) from OvHV-1, aligned with cellular and viral relatives. Grey shading indicates identity with the OvHV-1 sequence. Dashes indicate a gap in the alignment. For IL10, the greater variability in the N-terminal region maps to the predicted signal peptide sequences (underlined). For vFLIP, death effector domains (DED) predicted by ProSite are indicated. The ovine FLIP sequence is truncated at 220 amino acids.

A further interesting macavirus-specific gene is Ov1_2.7, which has 78% amino acid identity to a predicted ornithine decarboxylase (ODC) encoded by BoGHV6 (Bov2.b2), and 58% identity to ovine ODC. ODCs catalyse the first step in the biosynthesis of polyamines, such as spermine and spermidine, which are polycationic alkylamines with important roles in a variety of cellular functions [54]. ODCs were previously thought to be unique to BoGHV6 among herpesviruses and their role in herpesvirus biology remains unclear. However, as polyamines are able to enhance cellular proliferation and are highly enriched in tumour cells, it has been suggested that ODC might promote proliferation of quiescent cells infected by herpesviruses, and provide a more favourable environment for virus replication [19].

OvHV-1 encodes six further genes that are conserved in all sequenced macaviruses. Two of these, Ov1_4.5 and Ov1_9.0, encode putative homologues of the apoptosis regulatory factor Bcl-2 that are similar to orthologues in other macaviruses. Ov1_5.0 encodes a putative G-protein coupled receptor. Ov1_6.0 is a spliced gene with 3 exons, predicted to encode a transcription factor. Ov1_7.0 and Ov1_8.0 are both predicted to be viral glycoproteins, conserved with respect to BoGHV6 homologues but with lower sequence similarity with the positional orthologues in OvGHV2 and AlGHV1. Table 3 summarises the recognised macavirus-specific proteins in AlGHV1, OvGHV2, BoGHV6 and OvHV-1.

### OvHV-1 encodes a novel viral FLICE-like inhibitory protein (vFLIP)

Ov1_2.6 has 29% amino acid similarity with ovine FLICE-like inhibitory protein (cFLIP; also known as caspase-8 FADD-like apoptosis regulator (CFLAR)) (Fig. 4b), an important regulator of extrinsic (death-receptor-induced) apoptosis [55]. cFLIP encodes two death effector domains (DEDs), which mediate binding of caspase-8 and FADD to inhibit activation of the death-inducing signalling complex. cFLIP exists in long and short isoforms, and plays essential roles in immune regulation, cancer progression, and resistance to apoptosis. Homologues of cFLIP have not been described previously in macaviruses but are present in other gammaherpesviruses, including SaGHV2, Kaposi’s sarcoma herpesvirus (KSHV), EHV-2 and BoGHV4. These viral FLIP (vFLIP) proteins were first characterised by their ability to inhibit apoptosis, which, in common with the cellular cFLIP, is mediated through DEDs [56, 57]. Perhaps the best studied herpesviral vFLIP is the K13 (ORF71) protein of KSHV, which is expressed during viral latency and is a potent activator of NF-KB signalling, by which it promotes cellular proliferation and survival and contributes to viral persistence and oncogenesis [58].

While vFLIP proteins from other gammaherpesviruses typically have two DEDs, ProSite analysis predicted only a single DED in the C-terminal portion of Ov1_2.6 (Fig. 4b). However, it is possible that the N-terminal domain contains a second divergent DED as this region contains six helical regions and a RxDL motif, which are conserved elements in other DEDs (Fig. 4b). Although functional studies will be required to determine the activity of Ov1_2.6, the presence of a putative vFLIP in OvHV-1 is interesting because it appears to be unique among macaviruses. In addition, the presence of a vFLIP and two Bcl-2-like proteins (Ov1_4.5 and Ov1_9.0) in OvHV-1 suggests that this virus could regulate apoptosis through multiple distinct mechanisms, a property shared with KSHV, EHV-2 and SaGHV2.

### Defective orthologues of BoGHV6 proteins

Notably, four ORFs annotated in the BoGHV6 genome are present in OvHV-1 at the nucleotide level but the protein-coding sequences are truncated by the presence of one or more stop codons. These are Ov1_2.8 (Bov2.b3 in BoGHV6), Ov1_5.1 (Bv11.b1), Ov1_9.1 (Bov9.b1) and Ov1_9.2 (Bov9.b2). In order to determine whether these ORFs are similarly truncated in other isolates of OvHV-1, regions spanning these ORFs were amplified by PCR from tissues obtained from additional sheep, and from a sample of the original uncultured lung lavage cells from case JA548. Sanger sequencing of the PCR products showed that the ORFs for these genes were also disrupted in all isolates, confirming the accuracy of the genome assembly and that the interrupted structure of these genes is conserved across OvHV-1 isolates (Supplementary Figs. S3 - S5).

The presence of these shorter or defective ORFs suggests that these genes were lost following the divergence of OvHV-1 and BoGHV6. While the apparent loss of these four genes in OvHV-1 is interesting, it is important to note that no function for the predicted BoGHV6 proteins has yet been demonstrated. However, if Bov2.b3, Bv11.b1, Bov9.b1 and Bov9.b2 are indeed genuine viral proteins, it might be speculated that the loss of these functions within OvHV-1, and the loss of IL-10 in BoGHV6, could reflect adaptations to their respective hosts, or to differences in cell type specificity between the viruses during latency or lytic replication. Gammaherpesviruses typically exhibit viral latency in B or T lymphocyte subsets, although some virus species may infect cells of the monocyte/macrophage lineage (reviewed in [48]), whereas the lytic cycle generally occurs in epithelial cells to facilitate release of infectious virus into the environment.

Determination of the complete genome sequence of OvHV-1 has confirmed its close relationship to BoGHV6 and other macaviruses. However, additional work is needed to characterise the biological features of OvHV-1, including its cell and tissue tropism and disease association. The early work on OvHV-1 suggested that alveolar macrophages were the primary source of the virus [1, 2], although it was occasionally isolated from lymph nodes or other tissues [1, 2]. The close genetic relationship with BoGHV6, a lymphotropic herpesvirus, is therefore interesting as it suggests that lymphocytes might be an additional target for OvHV-1 *in vivo*. In this regard, a previous report described the detection of OvHV-1 in DNA from ovine peripheral blood mononuclear cells [59]. Interestingly, that study also found OvHV-1 in PBMC DNA from healthy cattle (54% positive) and bison (11%) indicating that OvHV-1 might infect multiple ruminant species. However, a subsequent analysis of lung tissue from 443 cattle from several European countries failed to identify any OvHV-1-positive animals by specific PCR [60]. Therefore, the range of host species and cell tropism of OvHV-1 remain to be further clarified. The availability of the complete viral genome sequence will facilitate such studies.

In recent years there has been growing interest in the diversity of macaviruses present in domestic and wild artiodactyl species, particularly with the aim of identifying novel MCF-associated viruses. Several groups have performed degenerate PCR using pan-herpesvirus primers in zoological collections to define the range of herpesviruses present [24, 25, 27, 28]. Those studies indicate a complex pattern of infection of viruses across species and it appears likely that additional uncharacterised macaviruses are circulating in artiodactyls. Notably, one of the short DNA polymerase gene fragments found in those studies is almost identical (175/176 nucleotide identity) to the corresponding region of OvHV-1. This sequence (GenBank AY237361) was reported as ‘domestic sheep-Lymphotropic Herpesvirus’ by Li and colleagues [27] and the close similarity suggests that this is likely to be OvHV-1. Notably, the source tissue for the isolate described by Li et al was peripheral blood leukocytes, again hinting that lymphocytes may be a site of latency for OvHV-1 *in vivo*.

The OvHV-1 genome has several extended regions that appear to be relatively poor in protein-coding genes, including much of the region to the left of Ov1_2.0, the region between ORF11 and ORF17, and the region between ORF69 and ORF73 (see Fig. 2). It is possible that these regions may yet encode proteins, perhaps from multiply spliced transcripts. Alternatively, these regions could encode non-protein coding RNAs such as microRNAs and long non-coding RNAs (lncRNA), as have been found in OvGHV2 and AlGHV1 and several other herpesviruses, [61, 62]. These macaviruses encode a major cluster of miRNAs between ORF11 and ORF17 [61, 63], while a further cluster is located close to the left-hand end of the genome. Some macavirus microRNAs have been shown to regulate expression of viral genes [64, 65], whereas others likely target host genes. While investigation of the microRNA-coding potential of OvHV-1 is outside the scope of the present study, it appears likely that future work will identify non-coding RNAs in this virus.

The annotation reported here is based on *in silico* ORF prediction and comparison to previously sequenced macavirus genomes. We have taken a conservative approach to annotating the coding regions, including only predicted ORFs supported by analogous genes in other gammaherpesviruses or their host species, and further work is required to analyse OvHV-1 transcription to confirm the structure of coding sequences and in particular spliced RNAs. Such experiments will require a more reproducible culture system for OvHV-1 than is currently available. Interestingly, recent studies using a range of high throughput methodologies have revealed that the transcriptional capability of herpesviruses is much more complex than previously appreciated, including for KSHV [66, 67], herpes simplex virus-1 [68] and Marek’s disease virus [69]. Similar detailed analysis may reveal greater transcriptional complexity for OvHV-1 and other macaviruses.

Finally, OvHV-1 is currently unclassified by the ICTV. In the early studies on OvHV-1 the virus was variously referred to as *Herpesvirus ovis* [9], Caprine herpesvirus-1 [13, 15], sheep pulmonary adenomatosis herpesvirus (SPAHV) [1], and bovid herpesvirus-4 [70]. Until the virus is formally classified by ICTV, we propose that this virus be provisionally designated ovine gammaherpesvirus type 1 (OvGHV1).

## Conflict of interest statement

The authors declare that there are no conflicts of interest.

## Acknowledgements

We thank the farmers who donated OPA-affected sheep for this study and the Moredun Bioservices Department for excellent animal care. We thank Richard Talbot and ARK Genomics (now Edinburgh Genomics) for Illumina sequencing and James Stewart and Laura Bianchessi for helpful discussions. This study was supported by funding from the Scottish Government Rural and Environment Science and Analytical Services Division (RESAS) to Moredun Research Institute and BioSS (BioSS-2-SAT).

## Notes

### Competing Interest Statement

The authors have declared no competing interest.

