## Supplementary figures and tables for "Genome sequence and annotation of ovine herpesvirus-1"

Paolo Ribeca, Patricia Dewar, Michelle L. McNab, Chris Cousens,
George C. Russell, David J. Griffiths[3]

**Supplementary Table S1 – Internal repeat regions in the OvHV-1 genome sequence.**

| Start | End | Length | Copy number | Total length | Sequence | Closest ORF |
|---|---|---|---|---|---|---|
| 922 | 1037 | 58 | 2 | 116 | CTACAAAAAGGGCCTTTCCACTATAACTTCTGCATAACCTGATTTATCACATGCTTAG | |
| 19336 | 19559 | 49 | 4.57 | 224 | CGTCTACCATGAGACTTAGTTTTGTGTTTTTGTGTTATGCTTAGGTGCG | Ov1_2.7 (ODC) |
| 49764 | 50353 | 27 | 21.85 | 590 | GGGCCTAGCTGCCCCCAGCAACCTACA | Ov1_5.1 |
| 135145 | 135521 | 18 | 20.94 | 377 | TCATCATCATCATCATCC | ORF73 |
| 135641 | 135747 | 36 | 2.972 | 107 | TACCACTAGGCTCACTTGCAGGAGTTTCTTCAGGGT | ORF73 |

**Supplementary Table S2 – Predicted spliced genes in macaviruses.**

| AlGHV1[1] | OvGHV2[1] | BoGHV6[1] | OvHV-1[1] | Function/comments |
|---|---|---|---|---|
| A2 (2) | Ov2 (2) | Bov2 (2) | Ov1_2.0 (2) | Basic leucine-zipper motif protein |
| | Ov2.5 (5) | | Ov1_2.5 (5) | Viral IL-10-like |
| ORF29 (2) | ORF29 (2) | ORF29 (2) | ORF29 (2) | DNA packaging terminase subunit |
| | ORF40 (2) | ORF40/41 (2) | ORF40 (2) | Helicase-primase complex |
| ORF50 (2) | ORF50 (2) | ORF50 (2) | ORF50 (2) | R-transactivator<br>Small 1st exon upstream of ORF49 |
| A6 (3) | Ov6 (3) | Bov6 (3) | Ov1_6.0 (3) | Putative transcription factor |
| | Ov8 (2) | Bov8 (2) | Ov1_8.0 (2) | Putative major envelope glycoprotein |
| ORF57 (2) | ORF57 (2) | ORF57 (2) | ORF57 (2) | Transcriptional control protein Mta |
| A9.5 (6) | Ov9.5 (6) | | | Potential IL-4 homologue |

[1] Number of exons is indicated in parentheses.

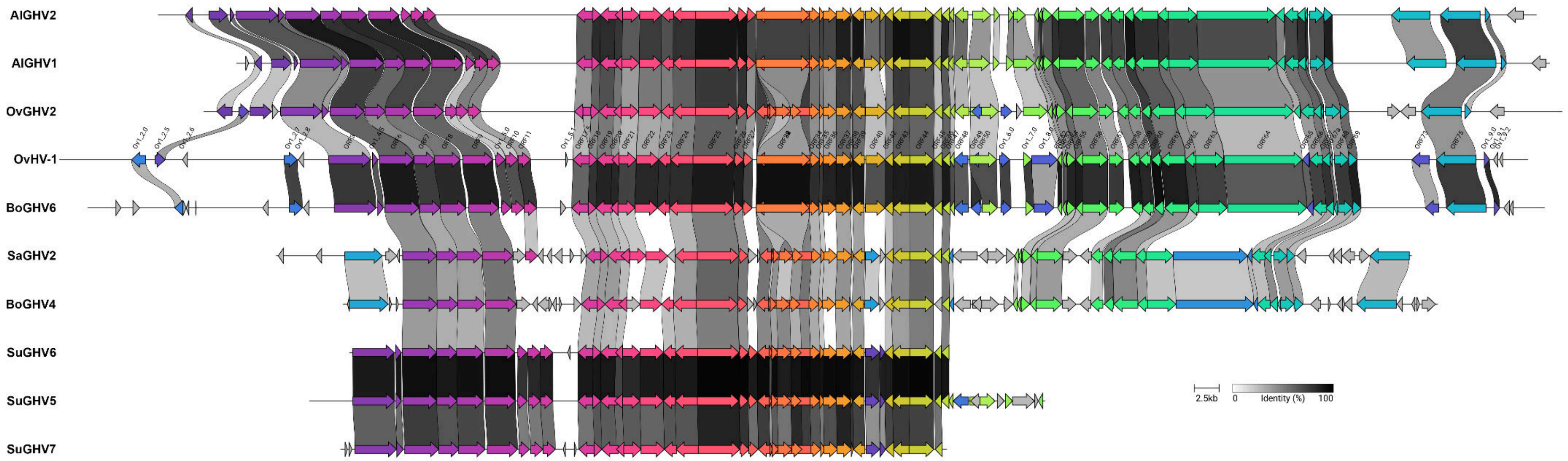


**Supplementary Figure S1 – Gene synteny between OvHV-1 and other herpesviruses of livestock.**
The figure shows a synteny plot for the genes in the viruses presented in the phylogenetic tree shown in Fig. 3a. This figure was generated with clinker [1]. Viruses are AlGHV2 (alcelaphine gammaherpesvirus 2; KF274499; nucleotides (nt) 1-137,090); AlGHV1 (alcelaphine gammaherpesvirus 1; NC_002531; nt 1-130,608); OvGHV2 (ovine gammaherpesvirus 2; AY839756; nt 1-135135); OvHV-1; (ovine herpesvirus 1; PV694339; 1-146035); BoGHV6 (bovine gammaherpesvirus 6; KJ705001; nt 1-144898); SaHV2 (saimiriine gammaherpesvirus 2 (herpesvirus saimiri)); X64346; 1-112930); BoGHV4 (bovine gammaherpesvirus 4 (partial); AF318573; nt 1-108873); SuGHV5 (suid gammaherpesvirus 5 (partial); AY170317; 1-59,673); SuGHV6 (suid gammaherpesvirus 6 (partial); AY478169; nt 1-73200); SuGHV7 (suid gammaherpesvirus 7 (partial); AY70316; nt 1-60326).

```
OvHV-1      MSLENSVAEEQMASEDEQQMSI---VEEEKKQKRKRKRICEESEERQAIIRARNKNASIK   57
BoGHV6      MSLQESTEEGQIAVKEEQQAAALA-EDPEKKQKRKRKRICEETEERQAIIRARNKNASIK   59
ALGHV1      -------------------MSQNSNSENPSPRKKRYVKMCDLTEEQKERRRSINRRASKN   41
ALGHV2      -------------------MFEYSSAPTQKLRKKKYVPMSELNEEQKEKRRIINRKASKN   41
OvGHV2      ------------------------MSDNKKPEKKTYPRMCDLTEEQRERRRSTNRRASRN   36
                                         . .*:    :.: .**::   *  *:.** :

OvHV-1      WREKNKKISETLTMVS | DVEKLEKENAQLHDTVYGLLREKHNLEFILEAHKATC-------   110
BoGHV6      WREKNKQIVETVT--R | DVEKLEKENAQLRDTVYGLLREKHNLEFILEAHKATC-------   110
ALGHV1      FLKRRRIFEEQ--QEK | GLINLKYENSRLRCQVEKRKDEIRILREWLNYHKCTTLQNYNTG   99
ALGHV2      FVQRRRLYEQQ--QEK | ELMDLKYENSRLRCQVEKNREELKQLRDWLCNHKYERAPNNNSE   99
OvGHV2      FKKRLQEHEQQ--QER | ELRDLIYTNMCLRGEIEKKKEEIRRLQYWLSTHNCLKLEGATPS   94
            : :: :   :          : .*   *  *:  :     * : *.  *  *:

OvHV-1      ------------KLEPAELFKLSAAQANTRSTVYCPTPQIKKHHQPISIQDQQQQHAKHR   158
BoGHV6      ------------KLEPVELFNLSAAKANTRSTVYCPTPQIQNHHQPISIQDHQQQHVKHR   158
ALGHV1      PPEPRVKVENSLEMQCATAFLN--------LDQQYTTNN-------LNIPETVSGNNTTN   144
ALGHV2      VSQLIENTANRCDLQNMNSLCM--------SSEQCVITN-------VNLLKPLPWTSPMD   144
OvGHV2      NSSSGG-----HGGEVTNDFYN--------SEQHLWSPQVGGSGCYFDPPAGSTWQMEDR   141
                          :    :                  :       ..

OvHV-1      GLPVIKQPQVIQAMPLHPQAVLHSLPLQPQQVIQAVSLQQPQVIRAVQVPQ-QVLQVQQN   217
BoGHV6      GLPVIKQPQVIQAMPLHQQAMVNSLPLQPQQVLQAVSLQQPQVVQALPVPQQQVLQVSQN   218
ALGHV1      GFAAA----------------TATLHTNCYE---KTL-----------------------   162
ALGHV2      GFAAT----------------AASFPTNFNS---GAT-----------------------   162
OvGHV2      GEGTA----------------SSSSP--FTG---EGS-----------------------   157
            *  .                   :

OvHV-1      QQRTVILPQNQIQFLPQLQIQIPSQQPAQQQQEIVPAHQAVPDVKQEQKTILDMSTNSVV   277
BoGHV6      QQNAMILQQNQIQFLPQLQIQIPAQQPVQQQPVF--TQQQQPVVKQEQKTLLDMSTNSVV   276
ALGHV1      -------ANNTNNFE------------AKLNCEVL------PSFTSALDDLLSIDWNNL-   196
ALGHV2      ---------AAVQAE------------VHSNNEVL------PSYTSTMDDFLLMD-----   190
OvGHV2      ---------------------------EQVLEELF------PETWLSVDISFDTELNML-   183
                                        :    .       *      .  :  .

OvHV-1      VKNEQDDSCSINIKREALSPVELKSNFRVPDNFPFEEDTLSDPEYFSNVIDSKFKDSVDE   337
BoGHV6      VKSEQGSSCALNIKTEAISPVDVKPNLRIPDDFPFEEDTLSDPEYFSNVMGSKLKEGVDD   336
ALGHV1      ------------------------------------------------------------   196
ALGHV2      ------------------------------------------------------------   190
OvGHV2      ------------------------------------------------------------   183

OvHV-1      GTESALSFLADIITAEAEKEAAPS--VSSSNNFQAPQPNTSHYSSSKVGSDWYHPTSAAT   395
BoGHV6      GAESALSFLANIVTAETEKEAALLPPVLSSSNQAALDALQGPQSNCYLSNAGYQLPLAAT   396
ALGHV1      ---------YNL------------------------------------------------   199
ALGHV2      ------------------------------------------------------------   190
OvGHV2      ---------HSL------------------------------------------------   186

OvHV-1      YNVNVEQPNKSTA-NSHVETDPWWKEGPQQAFNWLYE    431
BoGHV6      VGVNTEQQAKSATISQVEESDTLWKEGPQQAFNWLY-    432
ALGHV1      -------------------------------------    199
ALGHV2      -------------------------------------    190
OvGHV2      -------------------------------------    186
```

**Supplementary Figure S2.** CLUSTAL Omega alignment of predicted AlGHV1 A2 orthologues in OvGHV2, BoGHV6, AlGHV2 and OvHV-1. The AlGHV1 and OvGHV2 proteins are reported to be basic leucine zipper family transcription factors [2, 3]. Leucine residues are coloured red. Asterisks (*), colons (:), and periods (.) below the alignment indicate fully, strongly and weakly conserved amino acid residues, respectively. Dashes (-) indicate a gap in the alignment. The first and second exons of each protein are separated by a pipe symbol (|). Sequence similarity is evident in the N terminal region, providing support for the annotation of the OvHV-1 Ov1_2.0 protein as a spliced orthologue.

```
           M  K  L  V  L  A  I  A  V  M  C  S  I  F  C  A  S  T  L  V  P  *  P  S  Q
OvHV1     ATGAAGCTTGTTTTGGCAATAGCTGTCATGTGCAGCATTTTCTGTGCTTCTACTTTAGTTCCTTAACCTTCGCAG   75
JA542     ATGAAGCTTGTTTTGGCAATAGCTGTCATGTGCAGCATTTTCTGTGCTTCTACTTTAGTTCCTTAACCTTCGCAG   75
JA555     ATGAAGCTTGTTTTGGCAATAGCTGTCATGTGCAGCATTTTCTGTGCTTCTACTTTAGTTCCTTAACCTTCGCAG   75
JA502     ATGAAGCTTGTTTTGGCAATAGCTGTCATGTGCAGCATTTTCTGTGCTTCTACTTTAGTTCCTTAACCTTCGCAG   75
JSV32     ATGAAGCTTGTTTTGGCAATAGCTGTCATGTGCAGCATTTTCTGTGCTTCTACTTTAGTTCCTTAACCTTCGCAG   75
          ***************************************************************************

           L  S  G  L  L  R  Y  I  L  F  Q  V  D  T  F  I  N  G  L  C  F  N  L  N  C
OvHV1     CTATCTGGCCTTTTAAGATACATCCTTTTCCAGGTTGATACTTTTATCAACGGACTTTGTTTTAATTTAAACTGT   150
JA542     CTATCTGGCCTTTTAAGATACATCCTTTTCCAGGTTGATACTTTTATCAACGGACTTTGTTTTAATTTAAACTGT   150
JA555     CTATCTGGCCTTTTAAGATACATCCTTTTCCAGGTTGATACTTTTATCAACGGACTTTGTTTTAATTTAAACTGT   150
JA502     CTATCTGGCCTTTTAAGATACATCCTTTTCCAGGTTGATACTTTTATCAACGGACTTTGTTTTAATTTAAACTGT   150
JSV32     CTATCTGGCCTTTTAAGATACATCCTTTTCCAGGTTGATACTTTTATCAACGGACTTTGTTTTAATTTAAACTGT   150
          ***************************************************************************

           D  S  G  Y  G  A  N  I  F  D  N  L  H  L  P  T  H  I  P  K  C  F  S  G  K
OvHV1     GATAGTGGCTACGGAGCTAATATTTTTGATAATTTACACTTACCTACGCATATACCTAAATGCTTTAGCGGCAAG   225
JA542     GATAGTGGCTACGGAGCTAATATTTTTGATAATTTACACTTACCTACGCATATACCTAAATGCTTTAGCGGCAAG   225
JA555     GATAGTGGCTACGGAGCTAATATTTTTGATAATTTACACTTACCTACGCATATACCTAAATGCTTTAGCGGCAAG   225
JA502     GATAGTGGCTACGGAGCTAATATTTTTGATAATTTACACTTACCTACGCATATACCTAAATGCTTTAGCGGCAAG   225
JSV32     GATAGTGGCTACGGAGCTAATATTTTTGATAATTTACACTTACCTACGCATATACCTAAATGCTTTAGCGGCAAG   225
          ***************************************************************************

             (Y)
           F  N  K  T  S  C  L  K  W  S  S  V  N  I  A  T  Y  Y (Y) Y  L  N  Y  M  K
OvHV1     TTTAACAAGACAAGTTGCCTTAAGTGGTCTTCAGTCAACATTGCTACTTACTA---CTACCTCAACTACATGAAG   297
JA542     TTTTACAAGACAAGTTGCCTTAAGTGGTCTTCAGTCAACATTGCTACTTACTA---CTACCTCAACTACATGAAG   297
JA555     TTTAACAAGACAAGTTGCCTTAAGTGGTCTTCAGTCAACATTGCTACTTACTA---CTACCTCAACTACATGAAG   297
JA502     TTTAACAAGACAAGTTGCCTTAAGTGGTCTTCAGTCAACATTGCTACTTACTACTACTACCTCAACTACATGAAG   300
JSV32     TTTAACAAGACAAGTTGCCTTAAGTGGTCTTCAGTCAACATTGCTACTTACTA---CTACCTCAACTACATGAAG   297
          *** *************************************************   *******************

           P  D  S  D  V  Q  S  I  K  S  S  L  R  G  L  L  *  S  L  Q  Q  T  Y  P  T
OvHV1     CCAGATTCAGACGTACAAAGCATAAAAAGCAGCCTTAGGGGATTACTTTAAAGCCTTCAACAAACCTACCCAACT   357
JA542     CCAGATTCAGACGTACAAAGCATAAAAAGCAGCCTTAGGGGATTACTTTAAAGCCTTCAACAAACCTACCCAACT   357
JA555     CCAGATTCAGACGTACAAAGCATAAAAAGCAGCCTTAGGGGATTACTTTAAAGCCTTCAACAAACCTACCCAACT   357
JA502     CCAGATTCAGACGTACAAAGCATAAAAAGCAGCCTTAGGGGATTACTTTAAAGCCTTCAACAAACCTACCCAACT   360
JSV32     CCAGATTCAGACGTACAAAGCATAAAAAGCAGCCTTAGGGGATTACTTTAAAGCCTTCAACAAACCTACCCAACT   357
          ***************************************************************************

                                                              Y
           Q  T  Y  P  T  A  G  I  R  L  I  D  K  T  E  S  N  *  L  P  E  S  I  S  M
OvHV1     CAAACCTACCCAACTGCTGGAATTCGCTTAATTGACAAAACTGAATCAAACTAATTGCCTGAAAGCATCTCTATG   417
JA542     CAAACCTACCCAACTGCTGGAATTCGCTTAATTGACAAAACTGAATCAAACTAATTGCCTGAAAGCATCTCTATG   417
JA555     CAAACCTACCCAACTGCTGGAATTCGCTTAATTGACAAAACTGAATCAAACTAATTGCCTGAAAGCATCTCTATG   417
JA502     CAAACCTACCCAACTGCTGGAATTCGCTTAATTGACAAAACTGAATCAAACTACTTGCCTGAAAGCATCTCTATG   420
JSV32     CAAACCTACCCAACTGCTGGAATTCGCTTAATTGACAAAACTGAATCAAACTAATTGCCTGAAAGCATCTCTATG   417
          ***************************************************** *********************

                                                                             Q
           Q  T  Y  H  D  S  K  K  L  S  I  M  Q  G  L  R  G  L  I  Q  T  L  *  R  A
OvHV1     CAAACGTACCACGACAGCAAAAAGCTTTCTATAATGCAAGGTTTACGCGGCCTAATCCAAACACTTTAAAGAGCT   477
JA542     CAAACGTACCACGACAGCAAAAAGCTTTCTATAATGCAAGGTTTACGCGGCCTAATCCAAACACTTTAAAGAGCT   477
JA555     CAAACGTACCACGACAGCAAAAAGCTTTCTATAATGCAAGGTTTACGCGGCCTAATCCAAACACTTTAAAGAGCT   477
JA502     CAAACGTACCACGACAGCAAAAAGCTTTCTATAATGCAAGGTTTACGCGGCCTAATCCAAACACTTCAAAGAGCT   480
JSV32     CAAACGTACCACGACAGCAAAAAGCTTTCTATAATGCAAGGTTTACGCGGCCTAATCCAAACACTTTAAAGAGCT   477
          ***************************************************************************

           V  R  L  *
OvHV1     GTTAGACTTTAA  519
JA542     GTTAGACTTTAA  519
JA555     GTTAGACTTTAA  519
JA502     GTTAGACTTTAA  522
JSV32     GTTAGACTTTAA  519
          ************
```

**Supplementary Figure S3.** CLUSTAL Omega alignment of nucleotide sequence of Ov1_2.8 (BoGHV6 Bov2.b3 homologue) from five isolates of OvHV-1, showing the truncated reading frames. The deduced amino acid sequence is indicated above the alignment. Stop codons are marked in bold text. Asterisks below the alignment indicate conservation across all isolates. Dashes (-) indicate a gap in the alignment. Variable amino acids are highlighted in yellow. Note the presence of four internal stop codons in the reference sequence and in 3 other isolates, which truncates the protein to only 21 amino acids. Interestingly JA502 has only two internal stop codons, which raises the possibility that some isolates might have fully coding copies of this gene.

**a)**

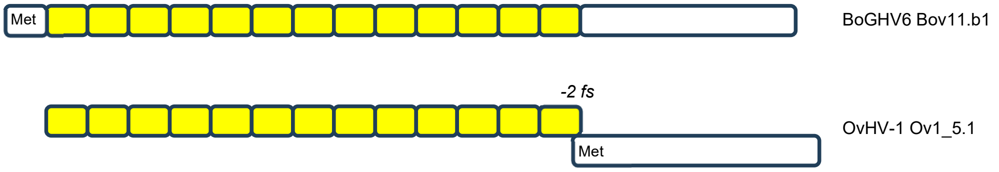


**b)**

```
BoGHV6 MPLTIVYEGLAAPSNLQGLAAPSNLQGLAAPSNLQGLAAPSNLQGLAAPSNLQGLAAPSN
OvHV-1 ------------------------------------------------------------

BoGHV6 LQGLAAPSNLQGLAAPSNLQGLAAPSNLQGLAAPSNLQGLAAPSNLQGLAAPSNLQGLAA
OvHV-1 ------------------------------------------------------------

BoGHV6 PSNLQGSAPKPEKSLLSAFENHTACQSTQKEAASYTFK--LINSHTVPKVRRKFKLAPTWTKVT
OvHV-1 MSNSQNSVPQLTESAPKAL-SPTACPKTQKPAAVTNFKQKLVNSHTVPKAQRKFKLAPTWTKLT
        ** * * *.  .*   *  . ***  *** **   **  *.******* .***********.*
```

**Supplementary Figure S4.** Comparison of BoGHV6 Bov11.b1 and OvHV-1 Ov1_5.1.

a) Representation of the structure of BoGHV6 Bov11.b1 and OvHV-1 Ov1_5.1. BoGHV6 Bov11.b1 has an N-terminal repeat region comprising 13 copies of the amino acid sequence LAAPSNLQG (represented by yellow boxes), which is in-frame with a unique (non-repetitive) C-terminal region (white box). In OvHV-1, the repeat and the unique regions are both present but are separated by a -2 frameshift (fs) immediately upstream of the methionine start codon. PCR and Sanger sequencing across this region in five OvHV-1 isolates found that the frameshift is conserved and that the number of upstream repeats varies between 20 and 33 copies (21 copies in the reference genome sequence. b) Amino acid sequence alignment of the two proteins. Repeat units are marked in alternating bold and underlined text.

**a)**

```
OvHV1   MTTKATTTEILLLGTEISLASTSLAPDNATDSPRASTSSNIYLEASLPPSGAPSHPPTER  60
JA542   MTTKATTTEILLLGTEISLASTSLAPDNATDSPRASTSSNIYLEASLPPSGAPSHPPTER  60
JA555   MTTKATTTEILLLGTEISLASTSLAPDNATDSPRASTSSNIYLEASLPPSGAPSHPPTER  60
JA502   MTTKATTTEILLLGTEISLASTSLAPDNATDSPRASTSSNIYLEASLPPSGAPSHPPTER  60
JA548   MTTKATTTEILLLGTEISLASTSLAPDNATDSPRASTSSNIYLEASLPPSGAPSHPPTER  60
JA564   MTTKATTTEILLLGTEISLASTSLAPDNATDSPRASTSSNIYLEASLPPSGAPSHPPTER  60
JSV32   MTTKATTTEILLLGTEISLASTSLAPDNATDSPRASTSSNIYLEASLPPSGAPSHPPTER  60
        ************************************************************

OvHV1   PHECPLPVETVPCNCKNEELLLYLLVLITLILLFCIIAIILFLILKIMRERDKRSRNRRD  120
JA542   PHECPLPVETVPCNCKNEELLLYLLVLITLILLFCIIAIILFLILKIMRERDKRSRNRRD  120
JA555   PHECPLPVETVPCNCKNEELLLYLLVLITLILLFCIIAIILFLILKIMRERDKRSRNRRD  120
JA502   PHECPLPVETVPCNCKNEELFLYLLVLITLILLFCIIAIILFLILKIMRERDKRSRNRRD  120
JA548   PHECPLPVETVPCNCKNEELLLYLLVLITLILLFCIIAIILFLILKIMRERDKRSRNRRD  120
JA564   PHECPLPVETVPCNCKNEELLLYLLVLITLILLFCIIAIILFLILKIMRERDKRSRNRRD  120
JSV32   PHECPLPVETVPCNCKNEELLLYLLVLITLILLFCIIAIILFLILKIMRERDKRSRNRRD  120
        ********************:***************************************

OvHV1   SDFSSGGSNSLKSPLKRDPSYMGLVF*  146
JA542   SDFSSGGSNSLKSPLKRDPSYMGLVF*  146
JA555   SDFSSGGSNSLKSPLKRDPSYMGLVF*  146
JA502   SDFSSGGSNSLKSPLKRDPSYMGLVF*  146
JA548   SDFSSGGSNSLKSPLKRDPSYMGLVF*  146
JA564   SDFSSGGSNSLKSPLKRDPSYMGLVF*  146
JSV32   SDFSSGGSNSLKSPLKRDPSYMGLVF*  146
        ***************************
```

**b)**

```
BoGHV6 MRCKFKACLNDQCMFNGYSIYFLETSHQTPTTTSVTTSAPTTTTFTSTLTTTTTTTTEPIVTTNATTNLLPRSTP  75
                                                                        TT +L   T
OvHV1  -----------------------------------------------------------MTTKATTTEILLLGTE  16

BoGHV6 NSLGSTSIAATNDASTTTNTTSSHGTVEVTQSSNATTVGLTTKGPHVIFEPPIKGTPCSCRERELFLWLLICCTC  150
        SL STS+A  N   +   +TSS+  +E +   +       T+ PH     P++  PC+C+  EL L+LL+  T
OvHV1  ISLASTSLAPDNATDSPRASTSSNIYLEASLPPSGAPSHPPTERPHEC-PLPVETVPCNCKNEELLLYLLVLITL  90

BoGHV6 LCIFLVILLLILLLLLKILRENRKPKTNPRRES----------------DAGYMQLVF*  191
       + +F +I +IL L+LKI+RE  K   N RR+S                D  YM LVF*
OvHV1  ILLFCIIAIILFLILKIMRERDKRSRN-RRDSDFSSGGSNSLKSPLKRDPSYMGLVF*  147
```

**c)**

```
OvHV-1                *FFIIGPNNKGGPPGCWPFKGCDQNHTFPLGTTINVSCTHDKNNSHNPNGRMWLNYSYSGN  60
                           GPN   GP  CWP   CD+N T PLG+ +N++C ++ +   NPNGRMWLNY+YSG+
BoGHV6 MVGSLCFQLYLILFLFLLFAGPNRVDGP--CWPNLKCDKNVTVPLGSVVNLTCKYNSSGLPNPNGRMWLNYTYSGS  74

OvHV-1 NGTLLLKYDNATYVNVTCSGPFDKKHGPLLCITWTNRKYLYL*  102
       NG+L LKY+N TY+NVTCSG FD+ HGPL+C+TWT+
BoGHV6 NGSLSLKYENHTYINVTCSGGFDQDHGPLVCLTWTSNSTYF*   115
```

**Supplementary Figure S5.** Confirmation of the sequence of OvHV-1 Ov1_9.1 and Ov1_9.2.

A region of the OvHV-1 genome spanning Ov1_9.0 to Ov1_9.2 was amplified by PCR from 6 independent isolates. Sequencing of the amplicons indicates that the sequences of the predicted Ov1_9.1 and Ov1_9.2 proteins are conserved and that both are truncated compared to the positional orthologues in BoGHV6. **(a)**. Alignment of predicted amino acid sequence of Ov1_9.1 from six isolates, including reamplification of JA548. **(b)**. BLASTP alignment of OvHV-1 Ov1_9.1 with BoGHV6 Bov9.1, indicating the extended region at the N terminus of the BoHV6 protein. **(c).** BLASTP alignment of OvHV-1 Ov1_9.2 with BoGHV6 Bov9.2, including the region upstream of the start methionine (green highlighting) to the closest upstream stop codon (*). Sequencing of this region from 6 isolates of OvHV-1 found no differences in the nucleotide sequence of Ov1_9.2 compared to the reference sequence.
